# Longitudinal Changes of Resting-State Functional Connectivity of Amygdala Following Fear Learning and Extinction

**DOI:** 10.1101/769034

**Authors:** Olga Martynova, Alina Tetereva, Vladislav Balaev, Galina Portnova, Vadim Ushakov, Alexey Ivanitsky

**Affiliations:** Institute of Higher Nervous Activity and Neurophysiology, Russian Academy of Sciences, 5a Butlerova Str., Moscow,117485, Russia; Centre for Cognition and Decision Making, National Research University Higher School of Economics, 20 Myasnitskaya Str., Moscow, 101000, Russia; Pushkin State Russian Language Institute, 6 Ak. Volgina Str., Moscow, 117485, Russia; National Research Center Kurchatov Institute, 1 Ak. Kurchatova Pl., Moscow, 123182, Russia

**Keywords:** fear conditioning, fear extinction, fMRI, resting state, functional connectivity, amygdala, PTSD

## Abstract

Altered functional connectivity of the amygdala has been observed in a resting state immediately after fear learning, even one day after aversive exposure. The persistence of increased resting-state functional connectivity (rsFC) of the amygdala has been a critical finding in patients with stress and anxiety disorders. However, longitudinal changes in amygdala rsFC have rarely been explored in healthy participants. To address this issue, we studied the rsFC of the amygdala in two groups of healthy volunteers. The control group participated in three fMRI scanning sessions of their resting state at the first visit, one day, and one week later. The experimental group participated in three fMRI sessions on the first day: a resting state before fear conditioning, a fear extinction session, and a resting state immediately after fear extinction. Furthermore, this group experienced scanning after one day and week. The fear-conditioning paradigm consisted of visual stimuli with a distinct rate of partial reinforcement by electric shock. During the extinction, we presented the same stimuli in another sequence without aversive pairing. In the control group, rsFC maps were statistically similar between sessions for the left and right amygdala. However, in the experimental group, the increased rsFC mainly of the left amygdala was observed after extinction, one day, and one week. The between-group comparison also demonstrated an increase in the left amygdala rsFC in the experimental group. Our results indicate that functional connections of the left amygdala influenced by fear learning may persist for several hours and days in the human brain.

## 1. Introduction

Although broader brain networks are involved in human fear processing than in animals, the amygdala was established as a hub of neural circuits for fear learning and extinction of fear memories (LeDoux, 2000; Etkin & Wager, 2009; Schumann et al., 2011). To date, most human studies have applied fear-conditioning protocols that allow the registering of the neural activation of brain regions in response to aversive stimulation by functional magnetic resonance imaging (fMRI). A variety of stimuli have been used with the conditioned stimulus (CS+), typically a visual cue or sound, which is always or partially reinforced by the unconditioned stimulus (US), a mild electrical shock, an aversive loud noise or, less frequently, an unpleasant smell or uncomfortable visceral stimulus (Gramsch et al., 2014; Kattoor et al., 2014). The inclusion of the second cue (CS−), which is not associated with the US, allows the comparison of neural and behavioral correlates of CS+ and CS−responses. Early fMRI studies found differential activity (to CS+ versus CS−) in the anterior cingulate cortex (ACC), anterior insula, hippocampus, and amygdala (Büchel et al., 1999). These findings have been replicated in fMRI (Rauch et al., 2006; Etkin & Wager, 2009; Shin & Liberzon, 2010; Andreatta et al., 2015) and magnetoencephalography (MEG) research (Balderston et al., 2014; Ma et al., 2013). However, a recent meta-analysis of fMRI showed inconsistency in the amygdala involvement in fear extinction learning in healthy individuals (Fullana et al., 2018) and the absence of amygdala activation in the late stage of fear learning (Fullana et al., 2016). Moreover, some studies reported the relatively poor reliability of amygdala activation (Nord et al., 2017), but good-to-excellent within-subject reproducibility of amygdala functional connectivity with the dorsomedial frontal/cingulate cortex in the emotional face-processing task (Nord et al., 2019).

Supporting the hypothesis that amygdala connectivity is a potential biomarker of stress-related maladaptation, an altered resting-state FC (rsFC) of the amygdala was also consistently reported in healthy groups after fear conditioning (Schultz et al., 2012), fear extinction, and fear reminder (Rauch et al., 2006; Feng et al., 2013; 2015). The increased rsFC of the amygdala with other regions of the fear network, such as the ventromedial prefrontal cortex (PFC), hippocampus, precuneus, and posterior cingulate cortex, or PCC), has also been found in patients with post-traumatic stress disorder (PTSD; Bluhm et al., 2009; Daniels et al., 2010; Liao et al., 2010; Dickie et al., 2011; Hayes et al., 2011; Rabinak et al., 2011; Yin et al., 2011; Zhou et al., 2012; Brown et al., 2014). Importantly, PTSD patients showed increased rsFC of the amygdala with cortical areas such as the ventromedial PFC compared with healthy controls in different fear learning protocols (Brown et al., 2014; Linnman et al., 2011). Trait anxiety was also associated with increasing coupling between the basolateral amygdala and anterior midcingulate cortex, supporting fear expression following extinction learning (Belleau et al., 2018). Remarkably, the rsFC of the amygdala might reflect distinctions in neural networks at the fear extinction stage in healthy participants (Feng et al., 2015) and might even predict the long-term expression of fear (Hermans et al., 2017).

Due to well-known hemispheric dominance and asymmetric interhemispheric information transfer in the human brain, the functional lateralization of the amygdala has also been studied regarding emotional learning. Most task-based fMRI studies reported the greater engagement of the left amygdala in emotional processing (Wright et al., 2001; Baas et al., 2004; Hardee et al., 2008; Hamann & Mao, 2002). The altered functional connectivity of the left amygdala has been found both in fear conditioning and post-task resting state activities in stress and anxiety disorders (Hahn et al., 2011; Prater et al., 2013; Baeken et al., 2014; Rus et al., 2017; Jung et al., 2018) that further support the hypothesis about differences in the functional role of the left and right amygdala in emotional regulation. Asymmetry in rsFC of amygdala was also shown in patients with social anxiety disorders (Yung et al., 2018). Simultaneously, the healthy population is also exposed to stressful events daily. This mild stress exposure could be stronger than aversive exposure during the classical fear-conditioning paradigm. Accordingly, we can assume that amygdala connectivity can also demonstrate longitude changes and fluctuations in healthy participants. However, there is a lack of findings describing long-term changes in the rsFC of lateral amygdala areas in a healthy population, which may serve as baseline data for comparison with anxiety and stress-related disorders.

The primary hypothesis of this study was that the amygdala’s neural activity established in fear learning might preserve a specific architecture or connections during a definite period after learning and fear memory extinction and could be observed in the resting state condition even in healthy participants with assorted anxiety levels. Three significant findings inspired this hypothesis: the altered rsFC of the amygdala in healthy subjects after fear learning (Shultz et al., 2012); the persistence of increased rsFC between the amygdala and hippocampus after several extinction and re-extinction training sessions (Hermans et al., 2017); and the altered rsFC of the amygdala in PTSD patients (Brown et al., 2014). Overall, findings about the amygdala’s rsFC suggest that post-learning neural activity may play a baseline role in memory reconsolidation. Our research questions concerned the extent to which the altered FC of the amygdala was preserved after a single session of fear learning with immediate extinction training. Additionally, we aimed to explore the possible asymmetry of long-term rsFC changes for the left and right amygdala in healthy participants. Thus, our study addresses two questions: how long the alterations in the amygdala’s FC during a resting state would persist a week after fear learning and extinction and would constitute a possible difference in the rsFC of the left and right amygdala depending on the fear learning or period. For this purpose, we compared rsFC of bilateral amygdalar regions with the whole brain in a control group of participants in three time points of a resting state (RS): at the first visit; in 24 hours; and seven days after the first RS session. Subsequently, we performed the same RS-scanning sessions for the experimental group that participated in the fear learning (FL) and fear extinction (FE) sessions, comparing the rsFC of the amygdala before, immediately after, 24 hours, and seven days after the FL session. We expected to show brain regions active in FE and explore differences in longitudinal changes of the rsFC of the amygdala in the control group and the experimental group with FL and FE.

## 2. Material and Methods

### 2.1. Participants

Two groups of healthy right-handed volunteers participated in the study. The control group consisted of 19 participants (18–32, mean age 26.2±3.89, five females). The experimental group consisted of 24 volunteers (18–30; 23.8±3.87, nine females). The difference in age was insignificant according to a Mann-Whitney U test (z = 1.93, p = 0.053) as well as in gender (z = 0.44, p = 0.66). All subjects had normal or corrected-to-normal vision and reported no history of psychiatric or neurological disorders. Additional exclusion criteria were the following: prior head trauma, any contraindications against MRI, medication affecting the central nervous system, consumption of drugs, excessive consumption of alcohol and nicotine, and pregnancy. Data of six additional participants were excluded from the analysis for the following reasons (1. malformation, 2. excessive head motions during MRI, and 3. apparatus failure). The study’s protocol was approved by the Ethical Committee of the Institute of Higher Nervous Activity and Neurophysiology of the Russian Academy of Science, according to the requirements of the Helsinki Declaration. All subjects provided written informed consent before the study.

### 2.2. Psychological Assessment and Analysis

All participants completed the State–Trait Anxiety Inventory (STAI; Spielberger et al., 1970) before each scanning session. Demographic characteristics and STAI scores for both groups of participants are reflected in Table 1. Nonparametric Wilcoxon Matched Pairs Test compared trait and state scores between sessions (days) and between groups with a Kruskal– Wallis test for independent samples.

### 2.3. Procedure

The control group participated in three resting state (RS) fMRI scanning sessions: first session (RS_0), a second session after 24 hours (RS_1), and the third session seven days after the first one (RS_7). Participants lay supine in an MRI scanner. The experimental group participated in five fMRI sessions: 1) RS_0 before FL; 2) Session with FE; 3) RS_FE immediately after FE; 4) RS_1 - in 1 day; and 5) RS_7 - a week after the FE session (Fig.1A). The duration of each RS and FE session was 10 minutes. There were five- to ten-minute breaks between sessions depending on the participant’s need, technical conditionings, and acquisition of structural MRI. During RS scanning sessions, participants were instructed to remain calm, with their eyes closed, to be awake, and to avoid purposely thinking about anything. We also stabilized participants’ heads with foam pads to diminish movement artifacts during MRI acquisition.

**Figure 1.**
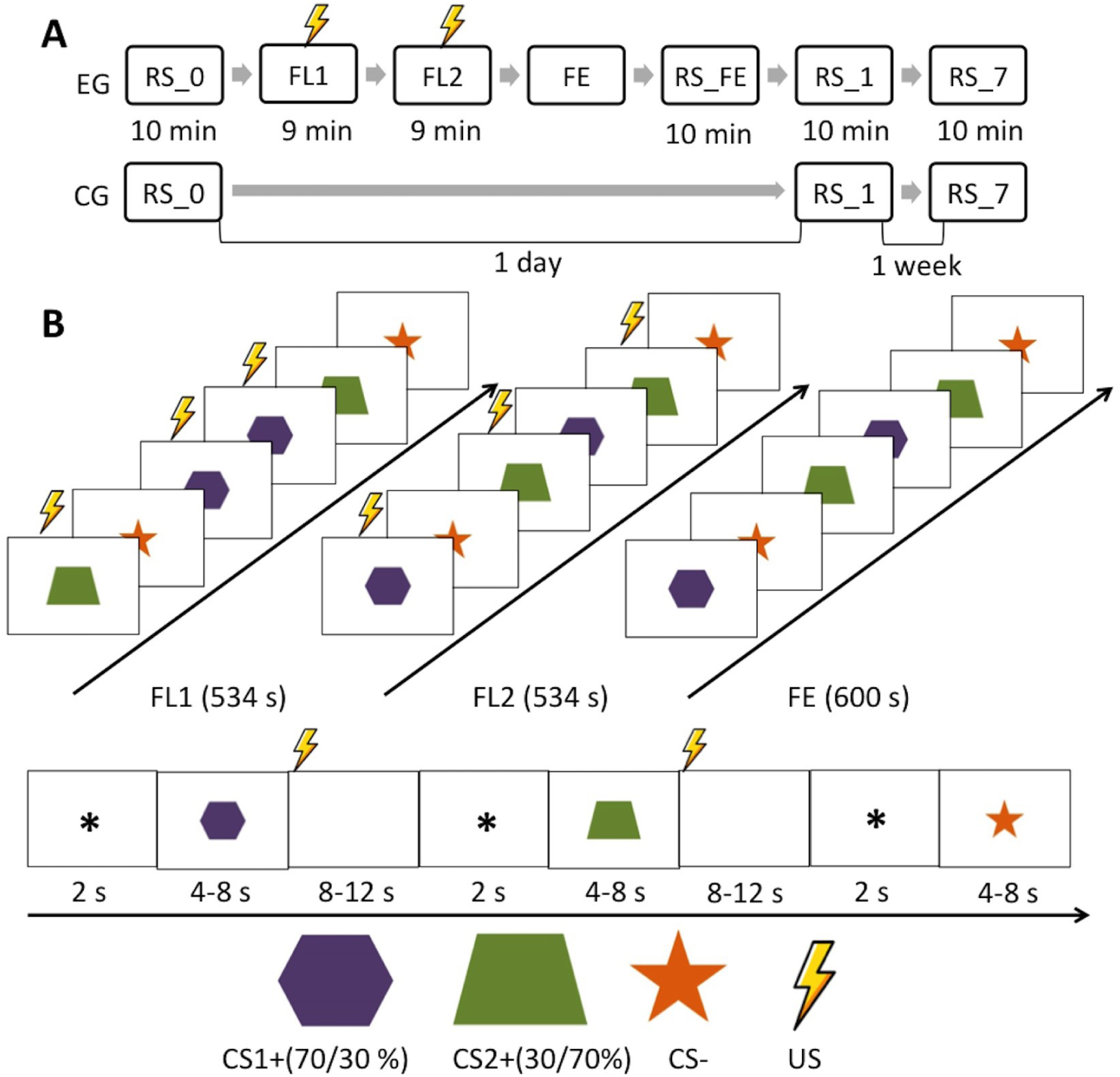
A- The study’s procedure for the control group (CG) and experimental group (EG) with fear learning: RS_0 – the first resting session, RS_1 – in one day, RS_7 – in seven days, FL1 and FL2 – two blocks of fear conditioning sessions, FE – fear extinction session, and RS_FE – resting state session after FE. B- Fear-conditioning paradigm with visual stimuli: CS1+ corresponds to the first type of stimuli with 70% reinforcement by electrical shock (US) in the first block of fear conditioning sessions (FL1), 30% reinforcement in the second block of fear conditioning sessions (FL2), and 0% reinforcement in the fear extinction session (FE); CS2+ corresponds to the second type of conditioned stimuli (30% in FL1, 70 % in FL2, and 0% in FE).

### 2.4. Fear Conditioning and Fear Extinction Paradigm

We hypothesized that fear acquisition would affect and cause alterations in amygdala connectivity, which has already been shown in previous studies (Shultz et al., 2012). We were more interested in the long-term effect of fear memories after FE, which was not studied so extensively, as it affects re-extinction training (Hermans et al., 2017) or the altered rsFC of the amygdala in PTSD patients. The other rationale to reduce the number of fMRI sessions (including imaging of fear learning) was the attempt to escape or to minimize an effect of context. The retention of fear memories could depend on the context, fMRI procedure, and even experimental environment (Lang et al., 2009). We tried to minimize the context effect by reducing the number of fMRI sessions on the first day, excluding scanning during and immediately after FL. Moreover, we ran two blocks of FL in a separate room of the behavioral lab, which was not adjoined to the MRI scanner facilities. We used a delay fear-conditioning paradigm with partial reinforcement, which consisted of three visual stimuli followed by a white screen: two conditioned stimuli (CS1+ and CS2+) were partially paired (30% and 70%) with US-a weak, short electrical shock, and one-stimulus CS−, which was never paired with US. All three types of stimuli were repeated 10 times. Before each stimulus, participants saw a fixation cross with a 2 s duration. The length of CS stimuli was randomly varied from 4 to 8 s, with 2 s increments. A white screen followed each CS presentation with a randomized duration from 8 to 12 s with 2 s increments. The average duration of the stimuli (CS1+, CS2+, and CS−) did not differ significantly within and between FL and FE sessions, according to a Kruskal-Wallis test. The US was administrated with no delay after the CS offset and coincided in time with the onset of the white screen. In the first block (FL1), the stimulus CS1+ was reinforced at the beginning of the white screen presentation with a 70% probability of electrical shock (US). CS2+ was reinforced with 30% probability, while SC-was never reinforced. In the second block (FL2), the reinforcement rate was 70% for CS2+ and 30% for CS1+. The blocks had a pseudorandom order of stimulus presentation (Table S1). The total duration of each FL block was 8 min, 54 s. During the FE session, we presented the same stimuli but in different pseudorandom order and longer sequence (10 min) without aversive reinforcement (Fig.1B). The FE session was conducted during fMRI. During the FE session, subjects were told to expect US but with a different reinforcement rule than in the two previous sessions. The US electrodes attached during the FE session were removed before the following resting-state session.

### 2.5. Conditioned Stimuli

During FL, the visual stimuli were shown using Presentation software (Albany, CA) on a monitor placed 50 cm from the participant, who was sitting on a comfortable chair in an isolated room with his or her right hand resting at the PC clipboard. During the FE session, the visual stimuli were presented via a video projector placed in the control room and a translucent screen and a mirror set up in the MRI room.

### 2.6. Unconditioned Stimulus

The US was 500 ms-duration electrical stimulation delivered via an AC (60 Hz) source (Contact Precision Instruments, Model SHK1, Boston, MA) through two surface cup electrodes (silver/silver chloride, 8 mm diameter, Biopac model EL258-RT, Goleta, CA). The electrodes were filled with electrolyte gel (Signa Gel, Parker Laboratories, Fairfield, NJ) and placed on the skin over the participant’s right tibial nerve above the right medial malleolus. Before FL, we performed a training session when we determined the maximum US intensity individually for each participant. The training session consisted of 10 presentations of electrical stimulation from a very low intensity of US to painful in two blocks with increasing and decreasing intensities of US. Each US presentation was rated by the subject on a scale from 0 to 10 (0 = no sensation, 10 = painful). In the first block, the intensity of the electrical stimulation was increased until the participant rated it as a 10, and in the second block, the intensity was gradually decreased until the participant rated it a 0. During FL sessions, the US intensity was set at the level averaged from two blocks at a level of 7 when each participant rated US as definitely painful, but tolerable.

### 2.7. Skin Conductance Recording and Analysis

To control FL, we recorded skin conductance responses (SCR). SCRs were measured using a direct current method (measurement voltage was 0.9 V) with a sampling rate of 4,000 Hz. Ag/AgCl electrodes (Medical Computer Systems LTD, Moscow, RF) with electrode gel were placed 14 mm apart on the left palm. The digitized signal was down-sampled using a low-pass filter of 16 Hz and manually cleared from movement artifacts in BrainVision Analyzer 2.1 (Brain Products, GmbH, Germany). The SCR for each CS trial was calculated by subtracting the mean skin conductance level measured 2 s before CS onset (during the fixation cross) from the highest skin conductance level recorded during the entire CS interval (Pineles et al., 2009). Values of SCRs were normalized for each participant, and then they were compared by nonparametric Wilcoxon Matched Pairs Test between CS1+ and CS−, CS2+ and CS−correspondingly in FL1, FL2, and FE sessions.

### 2.8. MRI Data Acquisition

MRI data were collected at the National Research Center Kurchatov Institute (Moscow, RF) with a 3T scanner (Magnetom Verio, Siemens, Germany) equipped with a 32-channel head coil. For each participant, sagittal high-resolution T1-weighted (anatomical) images were acquired using a T1 MP-RAGE sequence: TR 1470 ms, TE 1.76 ms, FA 9°, 176 slices with a slice thickness of 1 mm and a slice gap of 0.5 mm, and a 320 mm field of view with a matrix size of 320 x 320. The functional images were collected in the same sequence for both FE and RS sessions using a T2*-weighted echo planar imaging (EPI) sequence of 300 volumes with generalized autocalibrating partially parallel acquisition (GRAPPA), an acceleration factor equal to 4 (Preibisch et al., 2008), and the following sequence parameters: TR 2000 ms, TE 20 ms, FA 90°, 42 slices with a slice thickness of 2 mm and a slice gap of 0.6 mm, a field of view (FoV) of 200 mm, and an acquisition matrix of 98 × 98. To reduce the field inhomogeneities of EPI, we also acquired magnitude and phase images to apply a field mapping algorithm.

### 2.9. Functional MRI Analysis

The MRI data were preprocessed in the Statistical Parametric Mapping Version 8 (SPM8; Welcome Trust Centre for Neuroimaging, UK). Functional images were realigned with the mean functional image for motion correction. An excessive motion during scanning (more than 1.5 mm) was also an exclusion criterion for the further analysis of fMRI data. We compared averaged values of scan-to-scan movements in three dimensions between groups and sessions to ensure that there was no impact of head motions on the obtained results. Magnitude and phase values from field map images served for the calculation of the voxel displacement map (VDM) and the subsequent mapping to the mean functional image. Next, each fMRI volume was unwrapped using VDM. After co-registration of the mean functional image with the anatomical image, all functional images were normalized into the standard template of Montreal Neurological Institute (MNI) space with a voxel size of 1.5 × 1.5 × 1.5 mm in two stages by the SPM8 New Segment tool: segmentation of anatomical images into grey and white matter, cerebrospinal fluid, bones, and air, followed by the deformation of fMRI volumes and anatomical images using deformation fields. After the normalization procedure, we smoothed fMRI images using a Gaussian kernel with a full width at half maximum (FWHM) of 6 mm. Preprocessing pipelines were similar for task and rest fMRI data, except that we applied a fifth-order Butterworth band-pass filter to resting-state fMRI data with a frequency window from 0.01 Hz to 0.1 Hz.

During the final stage of preprocessing, we performed a regressing-out procedure to exclude motion patterns, breath-holding-induced signal changes (BHISC), and signals derived from ventricles from the fMRI data (Fox et al., 2009). BHISC was extracted as follows: first, we determined the times corresponding to the peaks from the respiration belt measurement. Subsequently, we convolved this “mask” vector with the respiration response function (Birn et al., 2008) and downsampled to match TR. The mask for ventricles was created in WFU PickAtlas 3.0.4 (Maldjian et al., 2003) and deformed to the individual space via the procedure described in the previous section. The resulting regression residuals were used for the further analysis of rsFC. The ventricles’ time series, six motion parameters, and BHISC were also applied as regressors in the general linear model analysis of the task session with a grey matter mask. At the segmentation stage, an individual gray matter image was converted to the binarized mask. The mask was applied to the same individual functional data for removing all the rest areas from rsFC and GLM analysis, except gray matter.

### 2.10. Functional Connectivity Analysis

We calculated the FC of the left and right amygdala with voxels from the whole brain separately. Masks of the amygdala were created for the left and right hemispheres in WFU PickAtlas 3.0.4 and coregistered with an individual anatomy image in the MNI space. The masks underwent inverse deformation to the individual subject space for each participant. The last step included the co-registration of the mask with the mean functional image. Furthermore, the mean BOLD signal was extracted for the left and right amygdala separately. Pearson correlation coefficients of the BOLD signal from the lateral amygdala seeds were calculated with a BOLD time series from every other voxel in the brain. Then, individual r statistics were normalized using a Fisher’s z transformation and subjected to a whole-brain one-sample t-test for all resting-state sessions. A paired t-test was applied for the comparison of RS correlation values between scanning sessions for both groups. A two-sample t-test was used for the comparison of rsFC differences in RS_0, RS_1, and RS_7 sessions between groups.

Additionally, we tested a relationship between individual values of the self-report scores for STAI with rsFC values for all scanning sessions of both groups, using normalized values in a Pearson correlation analysis. Following this, we ran separate multiple regressions in SPM8, testing scores of state/trait anxiety and depression as regressors and amygdala connectivity as the dependent variable. Depression scores were taken only on the first day and were subjected to correlation for all RS sessions, while STAI scores were taken repeatedly on days with scanning and used for correlation analysis with corresponding RS session data.

### 2.11. Analysis of Event-Related BOLD Response during Fear Extinction

We used the onset of each CS and the white screen (WS) presentation as the condition onset with 0 s duration and then convolved the event-related BOLD signal with the canonical hemodynamic response function (HRF) separately for CS1+, CS2+, CS−, WS after CS1+, WS after CS2+, and WS after CS−. To examine both the retention and the process of extinction of fear memories, we split the extinction session into the first half to test the retention effect and the second half to examine late extinction. According to this procedure, we estimated the BOLD dynamic derived the first and second half of the FE session.

After the convolution, the obtained vectors of the modeled BOLD response were used as regressors together with six motion parameters and a BOLD signal from ventricles (taken from the individual space) in the general linear model (GLM) for the first-level analysis for contrasts estimating event-related BOLD to WS against CS presentation by one-sample t-test. Altogether, we tested six contrasts—three contrasts for the retention and three for the extinction part of the FE session: (1) CS1+ >CS− minus WS after CS1+ and CS−, (2) CS2+ >CS− minus WS after CS2+ and CS−, and (3) (CS1+ plus CS2+) > CS− minus all WS. Because of the second-level analysis, we compared activation maps for the six contrasts and two additional contrasts: all WS versus all CS separately for the retention and extinction stages. We used a one-sample t-test for all contrasts and a paired t-test for the comparison of the first and second half of the FE session. Furthermore, using multiple regression models, we analyzed the dependence of the event-related BOLD response from STAI (State and Trait scores separately) and SCR for each participant.

The significance of the obtained statistical results for both rsFC and GLM analysis was estimated using a cluster size threshold adjusted for individual voxel type-I error with a voxel-wise cutoff of p<0.001 and cluster-size cutoff to get *p*_FWE_<0.05, corrected for multiple comparisons.

The lowest power or type-II error was observed in the pairwise comparison of the mean rsFC of amygdalar seeds in the control group. However, an estimation of power (1-type-II error) was higher than 90% for the average rsFC of the left amygdala in the following comparisons within the experimental group: RS_0 versus RS_FE and RS_0 versus RS_7. For the comparison of RS_0 and RS_1, the power was 0.70. Importantly, the power of the t-test comparing the left amygdala FC in RS_0 between groups was also higher than 0.95. Values of the left amygdala rsFC were less variable within subjects, keeping the standard deviation of the mean lower than 30%, while the right amygdala CCs changed more strongly, but with higher standard deviations (>70%).

## 3. Results

### 3.1. Comparison of Anxiety and Depression Scores between Sessions and Groups

All participants had scores for anxiety (STAI) within the moderate range (Table 1). There were no significant differences in state and trait scores among the three sessions for the control group. The trait scores were significantly lower for the first day than 24 hours (z=−2.38, p=0.02) and seven days for the experimental group (z=−2.98, p=2.97e-3). However, we did not observe significant distinctions in STAI scores between groups as between males and females.

### 3.2. Skin Conductance Changes during Fear Conditioning and Fear Extinction

Because we used two partially reinforced fear conditioning paradigms consisting of two blocks with a counterbalanced order of reinforcement, we observed the nonlinear dynamic of SCRs in response to CS in FL blocks and later in an FE session (Fig. 2, Fig. S1). SCRs increased significantly to CS1+ compared with CS−(z=−4.85, p=1.23e-6) and to CS2+ compared with CS−(z = − 3.10, p = 1.91e-3) in all FL blocks, while in FE sessions, SCR to CS−was lesser than CS1+, but higher than CS2+ (Fig.3): z = 2.39, p = 0.02, and z = − 2.73, p = 6.41e-3, correspondingly. Moreover, CS− was higher in the FE session than in two FL blocks (p = 6.04e-4). A trial-to-trial course of SCR level demonstrates uncertainty in the expectancy of US (Fig. S1), which also corresponds to the average accuracy rate of US prediction. Moreover, there were no significant variations in SCR level between a shorter or longer presentation of SC+.

**Figure 2.**
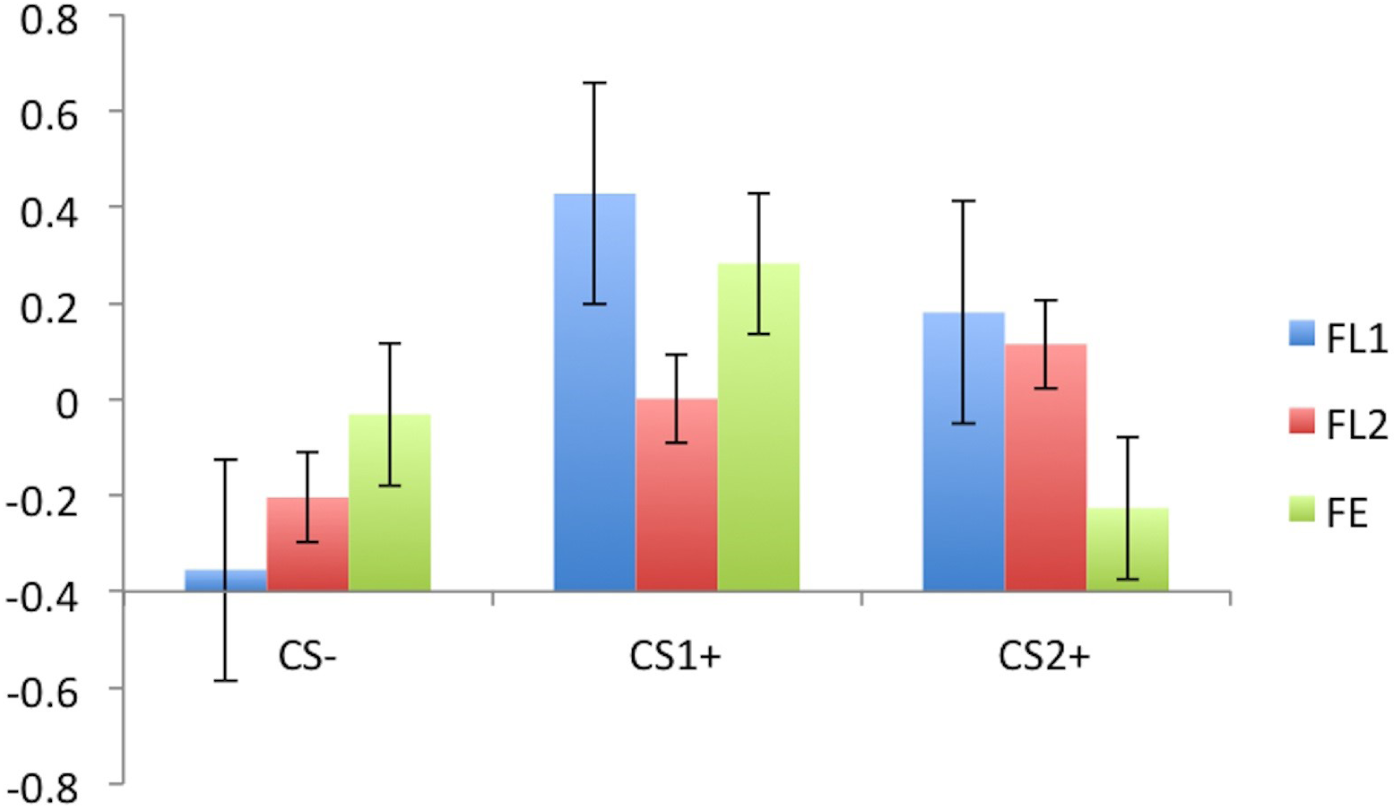
Averaged and normalized skin conductance responses (SCR) to conditioned stimuli (CS+) and stimuli without reinforcement by electrical shock (CS−): CS1+ corresponds to the first type of stimuli with 70% reinforcement in the first block of fear conditioning sessions (FL1), 30% reinforcement in the second block of fear conditioning sessions (FL2), and 0% reinforcement in the fear extinction session (FE); CS2+ corresponds to SCRs to the second type of conditioned stimuli (30% in FL1, 70 % in FL2, and 0% in FE).

After dividing SCR values during an FE session on two stages (retention and extinction), we observed the significant decrease of SCR to CS− comparing to CS1+ in both stages (z=−2.39, p=0.02, and z=−2.61, p=8.90e-3, correspondingly), while SCR to CS− and CS2+ did not differ significantly. There was also no difference between SCR to CS− in the first and second half of an FE session, and between SCR to CS1+. However, SCR to CS2+ decreased significantly in the second half of FE, compared with the first half (z=−2.67, p=7.60e-3).

### 3.3. Brain Activation During Fear Extinction

All examined comparisons of CS+ versus CS− at both stages of extinction provided activation maps only at an uncorrected level (p = 0.001), except for large clusters of activation in the occipital lobe. Only the joint analysis of contrasts corresponding to all WS versus all CS showed reliable results after correction for multiple comparisons. A one-sample t-test revealed an increased event-related BOLD response corresponding to the beginning of WS presentation after CS (p_FWE_ < 0.05, cluster size > 169) during the retention stage of an FE session in the bilateral areas of the occipital cortex (BA 18, 19), middle frontal gyri (BA 9,10), inferior frontal gyri (BA 44,47), and supramarginal gyri (BA 40; Table S2A, Fig. 3). During the extinction stage (the second part of an FE session), we observed an increase in the BOLD response to WS after CS (p_FWE_ < 0.05, cluster size > 169) in the left lingual gyrus (BA 18, 19), parahippocampal gyrus (BA 30), left medial frontal gyrus (BA 10), and right precuneus (BA 31; Table S2B, Fig. 3). However, we observed no significant correlation of the BOLD response with SCR values.

**Figure 3.**
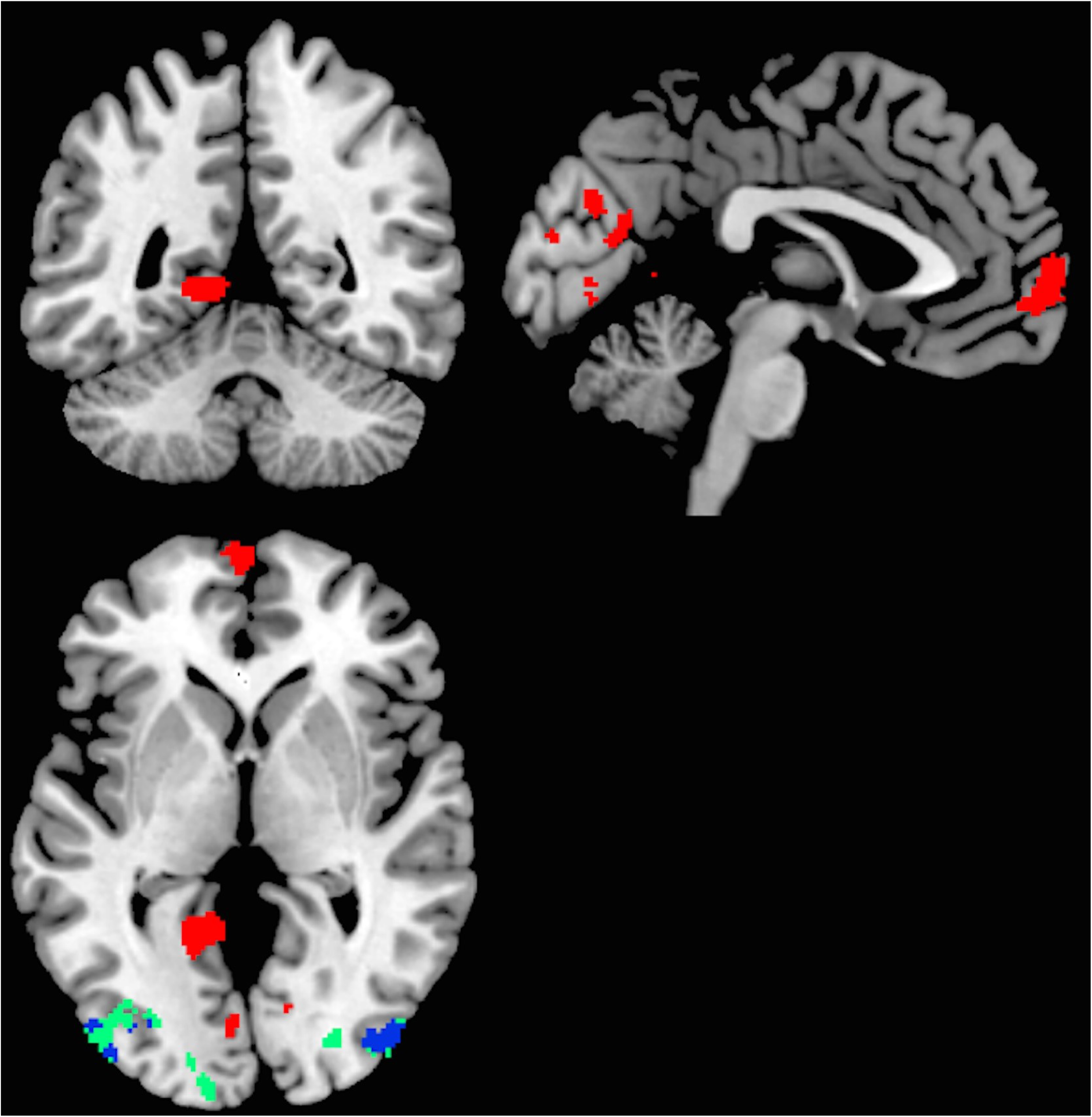
The event-related BOLD response in the contrast of the white screen after conditioned stimuli in the retention stage of fear memory (blue color corresponds to positive response) and extinction stage (red – positive, green – negative response) in the experimental group (cluster size threshold > 169, P_FWE_<0.05).

### 3.4. Functional Connectivity in Resting State Sessions 24 Hours and One Week after Fear Extinction

In the control group, rsFC maps were statistically similar between all RS sessions for both amygdala seeds. In the experimental group, we obtained a significant increase of the left amygdala rsFC in the session after FE (RS_FE), compared to the RS_0 session before fear conditioning with the following areas (Fig. 4A, Table S3A): bilateral precuneus, right supplementary motor area and left middle and superior frontal gyri, middle temporal gyrus, postcentral gyrus, superior temporal pole, and midcingulate area (P_FWE_<0.05). One day after FE, we observed increased connectivity of the left amygdala with the right temporal gyrus, right cuneus, and left precentral gyrus (Fig 4B, Table S3B). Remarkably, rsFC of the left amygdala was also significantly increased in the RS_7 session compared to the RS_0 (Fig. 4C, Table S3C) with the following areas: bilateral temporal and superior frontal gyri, right superior temporal cortex and calcarine sulcus, right cingulate cortex, and midcingulate area.

**Figure 4.**
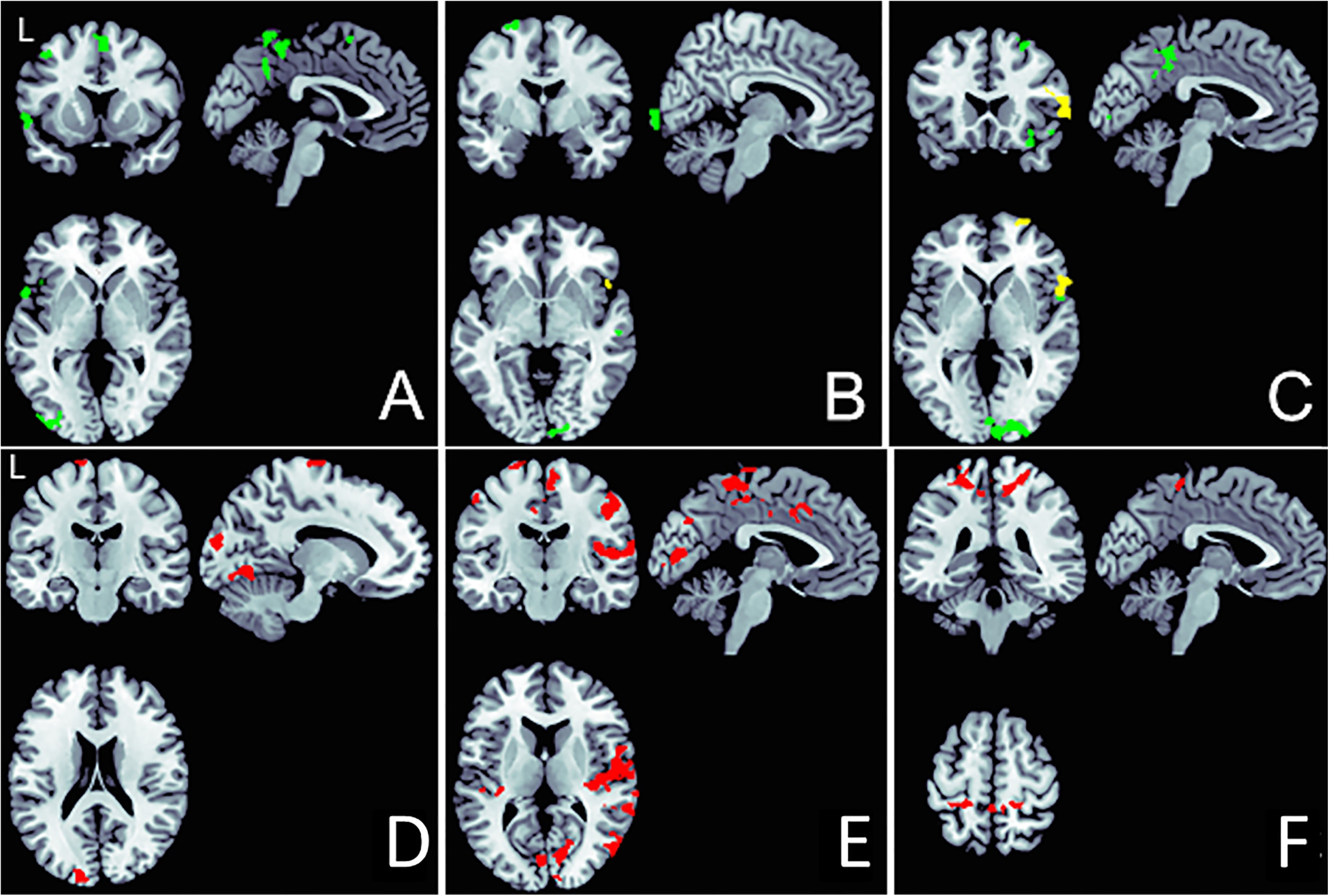
Brain areas showed significant functional connectivity with the left amygdala (in green) and right amygdala (in yellow): A-after fear extinction, MNI coordinates: x = −2, y= 12, z = 0; B - in 24 hours, x = 6, y = −2, z = −4; and C - in 7 days, x = −4, y = 22, z = 1; cluster size threshold >178, P_FWE_ < 0.05. Brain areas showed significant functional connectivity (FC) in the experimental group, compared with the control group: D – FC with the left amygdala in 24 hours, MNI coordinates: x = −13, y = −16, z = 20; E – FC with the left amygdala seven days after the first session, x = −4, y = −16, z = 10, and F – FC with the right amygdala seven days after the first session, x = −4, y = −36, and z = 61 (cluster size threshold > 213, P_FWE_<0.05).

There was no difference between rsFC of the right amygdala between RS_0 and RS_FE sessions. One day after FE, the right amygdala showed an increase in rsFC with the right inferior frontal gyrus (Fig. 4B, Table S3B) and expressed increased rsFC with the middle frontal gyrus, the opercular part of the inferior frontal gyrus, and the left superior parietal lobule seven days after FE compared to the first session before experimental exposure (Fig. 4C, Table S3C).

The between-group comparison also showed a significant difference only for the left amygdala rsFC between RS_0 and RS_1 sessions (cluster threshold > 217 voxels, PFWE<0.05). For the experimental group, the left amygdala rsFC with left frontal gyrus was increased, compared with the control group in the first resting state session (RS_0; Table S4A). On the second day (RS_1), the left amygdala’s rsFC was higher with the left lingual gyrus, paracentral lobule, and superior occipital gyrus in the experimental group than in the control group (Fig. 4D, Table S4B). In a one-week session, we observed even more significant variations between the control and experimental group. The two-sample t-test showed a significantly higher rsFC of the left amygdala with a bilateral precentral gyri, superior temporal gyri and calcarine sulcus, left paracentral lobule, right insula, midcingulate area, middle temporal gyrus, and left cuneus (Fig. 4E, Table S4C) in the experimental group. The right amygdala’s rsFC was greater only than the right paracentral lobule in the experimental group, compared to control group rsFC data (Fig. 4F, Table S4C).

We examined possible conjunctions between areas exhibiting activity during FE and changes between sessions in functional coupling with the amygdala. The masks of clusters with a significant increase of BOLD response overlapped with masks of rsFC. The area of increased coupling of the left amygdala with the left superior frontal gyrus (BA 6) overlapped in several within-session comparisons (Fig. 5A, Table S5). We also found one area in the right inferior frontal gyrus (BA 47) in the observed BOLD response in the retention stage and in within-session differences in rsFC with the left and right amygdala (Fig. 5B,Table S5).

**Figure 5.**
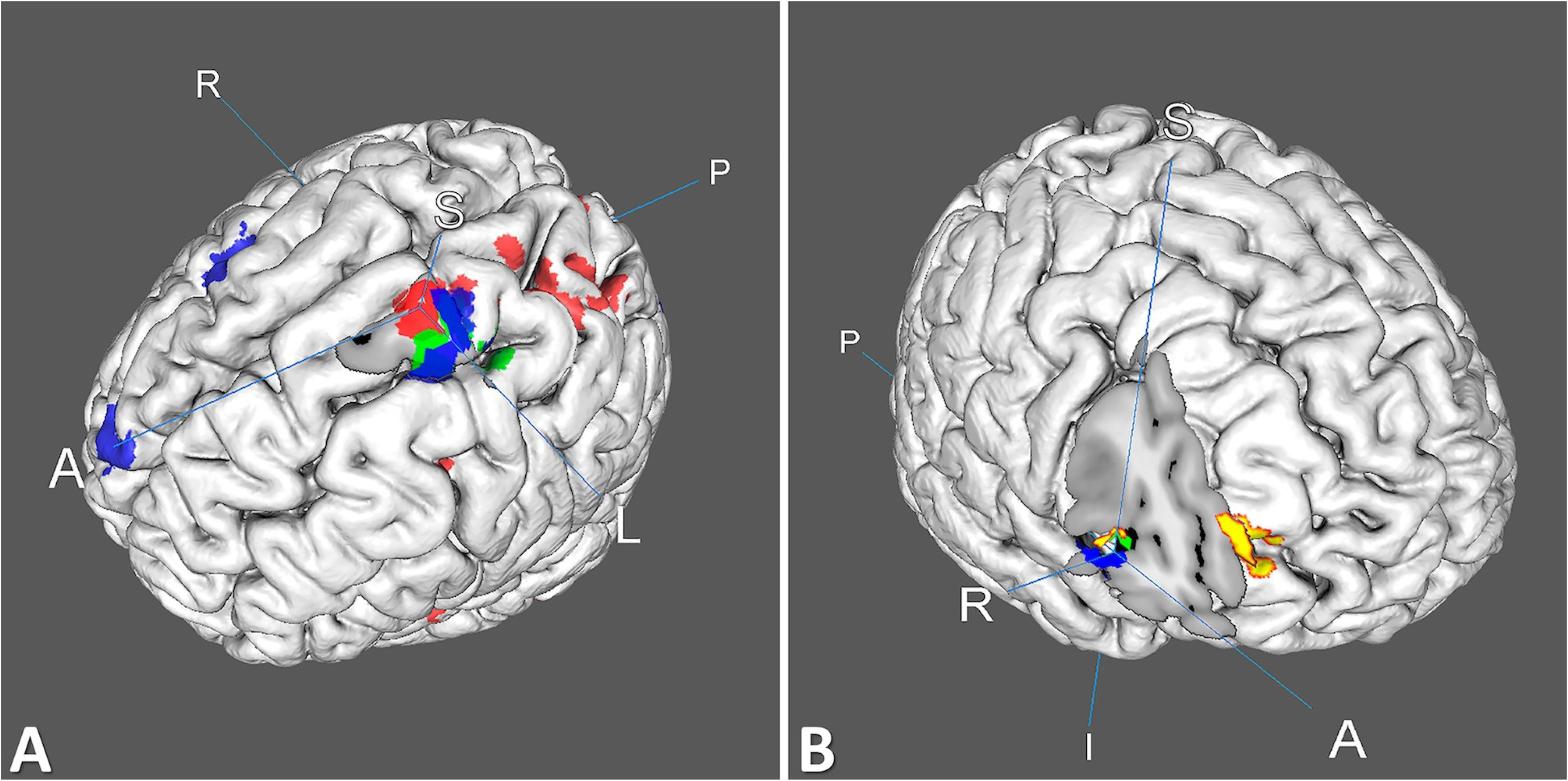
(A) The spatial overlap of areas with differences in the FC of the left amygdala in RS sessions: red color corresponds to the comparison of RS_FE and RS_0, green – to RS_1 and RS_0, blue – to RS_7 and RS_0. (B) The spatial overlap of areas with variations in the FC of the right and left amygdala and event-related BOLD response in the retention stage: blue color corresponds to comparisons of RS_7 and FE for the left amygdala, green – to RS_1 and FE for the right amygdala FC, and yellow – to RS_7 and FE for the right amygdala.

### 3.5. Correlation of BOLD Response and Functional Connectivity of Amygdala with Anxiety Scores

During an FE session, the event-related BOLD response to WS versus CS showed a significant association with trait anxiety scores in the orbital part of the right middle frontal gyrus (BA11) and the inferior frontal gyrus, *pars triangularis* (Table S6).

In the experimental group, we revealed a significant relationship between the strength of the right amygdala’s rsFC with the medial frontal gyrus and trait anxiety scores only for the first RS session (RS0) in the experimental group (Table S6). However, in the control group, we found significant correlations between trait scores and the left amygdala’s rsFC with the right hippocampus in RS1, as well as between-state scores and the right amygdala’s rsFC with the right precuneus and left middle frontal gyrus in RS7 (Table S6).

## 4. Discussion

In this study, we compared daily and weekly changes of rsFC of the left and right amygdala in two groups of healthy participants: the control group and experimental group, which underwent a fear learning procedure and the next, short extinction training in the partially reinforced fear conditioning protocol. Our findings indicate that plastic changes of the amygdala functional connectivity influenced by fear conditioning and subsequent extinction may be preserved in the RS condition after several hours and days in the healthy human brain. Importantly, altered rsFC after fear learning was more prominent for the left than the right amygdala. In particular, we observed increased rsFC only for the left amygdala with several brain areas, including dorsal PFC and the midcingulate area, immediately after extinction training in the experimental group, compared with a baseline (RS session before fear learning). After 24 hours, we still observed altered rsFC for the left amygdala but with fewer brain regions, while in seven days, we found increased rsFC of the left amygdala with more areas, including the dorsal PFC, the right cingulate cortex, and the midcingulate area. The rsFC of the right amygdala with dorsolateral PFC also increased in seven days to a greater extent than in RS in 24 hours, compared with the first RS session.

Our findings are partially consistent with previous results on the altered rsFC of the amygdala after fear learning and extinction (Rauch et al., 2006; Feng et al., 2013; 2015; Hermans et al., 2017). Notably, the amygdala’s rsFC with dorsal PFC stayed altered after fear learning (Schulz et al., 2012) and in 24 hours, but returned to the baseline level one week after fear conditioning in healthy participants (Schultz et al., 2014). Schultz et al. (2012) used the full reinforcement paradigm and rsFC of the averaged left and right amygdala. In our study, we employed partial reinforcement fear conditioning protocol and observed the increase of rsFC in 24 hours and even seven days after fear learning, specifically for the left amygdala. The regions showed a persistent increase in the functional coupling with the left amygdala included not only the dorsal PFC, which was previously suggested to reflect long-term memory retrieval (Stock et al., 2009), but also the cingulate and midcingulate cortex. The increased rsFC between the amygdala and cingulate cortex was also observed immediately after fear learning (Feng et al., 2013), was shown in PTSD patients (Dickie et al., 2011; Rabinak et al., 2011; Yin et al., 2011; Zhou et al., 2012; Brown et al., 2014), and emerged early in animal and human research of PTSD, suggesting the cingulate cortex played an inhibiting role over amygdala activity during fear memory extinction (Gilboa et al., 2004). We did not find a long-term alteration of amygdala rsFC with the ventromedial PFC and hippocampus, as was shown in previous studies (Hayes et al., 2011; Feng et al., 2013; Hermans et al., 2017). Instead, our results imply the persistently increased coupling of the left amygdala with dorsal PFC, midcingulate cortex, middle and superior temporal cortex, precentral and postcentral gyri, and cuneus after fear learning. We did not use pre-defined or hypothesis-driven regions of interest (ROI) like the aforementioned studies, but analyses of the whole-brain rsFC with bilateral amygdala seeds. The latter could explain the absence of a substantial increase of rsFC of the amygdala with the medial PFC and hippocampus, frequently reported early in PTSD and anxiety patients (Dickie et al., 2011; Brown et al., 2014). It is also possible that altered rsFC with these brain areas is more prominent in the clinical population than in healthy participants.

After overlapping masks of significant clusters from the paired comparison of amygdala rsFC between different scanning sessions, we found only one brain area in the dorsal PFC (left superior frontal gyrus, BA 6), where masks of left amygdala coupling overlapped or adjoined. This area of increased FC with the left amygdala appeared when comparing the first session prior FL with sessions after FE, one day and one week after FL and FE. The other region of overlap between BOLD responses in the retention stage with longitude changes of resting-state FC of both lateral amygdala seeds was observed in the dorsolateral PFC (right frontal inferior gyrus, BA47). The consistent activation of the dorsolateral PFC was mentioned early in the meta-analysis of FE (Fullana et al., 2018). The other study regarding the context effect on FE also reported the enhanced activation of dorsolateral PFC both during extinction and renewal conditions when subjects returned to the fear-learning context (Icenhour et al., 2015). Compared with the medial PFC, the dorsal and dorsolateral cortex is an infrequent target or ROI in resting-state research of amygdala connectivity. We found only one study concerning prefrontal– amygdala connectivity during fear extinction recall in adolescents reported significant negative FC between the dorsolateral PFC and the amygdala, which was positively correlated with state anxiety (Ganella et al., 2017). Our results suggest that plastic changes between the amygdala and dorsolateral PFC formed during fear learning and extinction, which can also be depicted in post-learning resting-state FC.

To support persistent long-term changes of amygdala connectivity after fear learning, we did not find significant alterations in the amygdala’s rsFC in the control group between three RS sessions (baseline, in 24 hours, and in one week), while the comparison of the rsFC between the control and experimental group showed differences in the rsFC of the left amygdala. The rsFC of the left amygdala with several brain regions was higher in the experimental group in 24 hours and seven days after the first RS session compared with the control group. The significant differences between groups occurred in one week and included increased rsFC of the left amygdala with bilateral precentral gyri, superior temporal gyri and calcarine sulcus, left paracentral lobule, right insula, middle temporal gyrus, cuneus, and midcingulate area in the experimental group. The right amygdala’s rsFC was greater only with the right paracentral lobule in the experimental group, compared to the control group. The functional lateralization of the amygdala has been discussed early concerning emotional regulation and the processing of threats (Phan et al., 2002; Costafreda et al., 2008; Baeken et al., 2014). Task-based fMRI studies have reported that the left amygdala was more active in emotional learning tasks (Phelps et al., 2001; Wright et al., 2001; Baas et al., 2004; Yun et al., 2017). More frequently, studies on anxiety observed the altered FC of the left amygdala (Hahn et al., 2011; Prater et al., 2013; Rus et al., 2017; Jung et al., 2018). Consequently, our results suggest that the left amygdala may play a more sustained role in long-term memory retention than the right amygdala, even in the case of healthy participants.

Notably, observed changes in the left amygdala’s rsFC could be affected by several factors. First, we used the partially reinforced fear conditioning protocol in the present study. Emotional responses measured by SCR indicate that participants learned to expect an aversive outcome after CS1+ and CS2+ during the fear learning session. However, both the retention and extinction stages SCR to CS−, which were not reinforced in FL, increased and even exceeded SCR to one of CS+. This result could occur due to the uncertainty of aversive outcomes for three conditioned stimuli during an FE session, as participants were not aware of the new order of reinforcement. When we separately analyzed SCR in the first and second half of the FE session, we found that fear memories concerning CS1+ were more resistant to extinction than CS2+, which also supports previous observations that uncertain reinforcement influences and diminishes the extinction of fear (Milad et al., 2009; Schiller et al., 2013; Feng et al., 2015). The observed significant alterations of amygdala FC in the resting state after FL and FE in the experimental group may imply some retention of fear memories due to the uncertainty-dependent effect of partial reinforcement. Moreover, we assume the random order of CS presentation, random duration of stimuli, and white screens between them in the fear learning procedure might strengthen the uncertainty and bring the fear learning procedure closer to the stress modeling paradigm.

Second, in the case of human studies, it is impossible to separate an effect on context and the spontaneous recovery of fear memory, as in the case of animal studies (Rescorla, 2004). We cannot entirely exclude the influence of context (the same experimental environment) on the long-term changes in the rsFC of the amygdala. However, these changes were more prominent for the left amygdala, which further supports the functional asymmetry of the bilateral amygdala in the sustained processing of fear memories. Third, we assume a specific role of the individual anxiety level and possible weekly changes in the personal situation of participants. Jung et al. (2018) have reported about the altered rsFC of the left amygdala in patients with social anxiety disorder. However, in our study, only the right amygdala’s rsFC with the right medial frontal gyrus correlated significantly with trait anxiety scores in the first baseline RS session in the experimental group. While there were no significant variations between group anxiety scores, we found an association between trait and state anxiety and rsFC of the amygdala only in the control group on the second day and one week after the first session. This finding indicates an effect of strong individual variability of human rsFC, which may diminish with an enlarged study sample. However, the absence of a reliable correlation or rsFC with state anxiety in the experimental group in 24 hours and one week after FL strongly suggest the observed persistence in the rsFC of the left amygdala is associated with the long-term expression of fear memory rather than with the effect of context regarding participants’ anxiety level.

Finally, the event-related BOLD response during the retention stage of the FE session occurred significantly in the bilateral areas of the visual cortex and dorsolateral PFC, while in the extinction stage, we observed an increase of fMRI activation in the right precuneus, left visual cortex, parahippocampal gyrus, and medial PFC. However, the amygdalar complex did not show specific activation, even at an uncorrected level. Therefore, our findings are only partially consistent with previous data, as many fMRI studies found a fear response in the amygdala, ACC, anterior insula, and hippocampus (Andreatta et al., 2015; Zhou et al., 2012). Activation in the amygdala, anterior insula, ACC, and left PFC have also been observed while subjects viewed a movie of someone else undergoing fear conditioning (Ma et al., 2013), suggesting similar processes of fear learning can be elicited without directly experiencing the US. Over the course of fear conditioning, amygdala activation typically decreases, as does hippocampal activation during trace conditioning (Reinhardt et al., 2010). However, a recent meta-analysis of fear conditioning experiments did not identify consistent amygdala involvement during fear conditioning (Fullana et al., 2016) and extinction (Fullana et al., 2018) in healthy participants, but instead confirmed large-scale activations in the cingulate cortex, medial PFC, and anterior insula, as well as additional cortical regions (supplementary motor area, dorsolateral PFC, and precuneus), ventral striatum, and midbrain. Under the comparison of an extinct threat stimulus with a non-extinct threat stimulus, more consistent activation was observed in the dorsolateral and ventromedial PFC, together with subcortical areas including the hippocampus (Fullana et al., 2018). The prevailing opinion, based on findings in animal research, postulates the amygdala is involved with the monitoring of external threats and learning how to escape them (LeDoux, 2014). However, a mild exposure to aversive stimuli in human fear conditioning protocols is very far from real threatening events that could explain inconsistent data on amygdala activation during fear conditioning in healthy participants. The other possible explanation is related to the limited spatial resolution of conventional fMRI due to both experimental design and analysis pipelines requiring averaging across trials, subjects, and strict correction for multiple comparisons. Because even in healthy controls, individual variability in response to fear conditioning is very high, we suggest using multivariate fMRI or machine learning approaches to analyze individual responses to emotional stimuli for further studies.

One of the study’s limitations concerns the sample size. Overall, the analysis of power of sample size confirms significant alterations of left amygdala connectivity within the experimental group (type II error > 95%). However, the absence of longitude changes in FC in the control group should be confirmed by collecting data from more participants. The other limitation and perspective for the further research of asymmetry in amygdala connectivity is the use of the amygdala’s subregions as basolateral and centromedial amygdala instead of extracting signal from the whole amygdala. Subregions of the amygdala are differently involved in the fear conditioning process (Laurent et al., 2008) and show disparate connectivity patterns (Belleau et al., 2018).

## 5. Conclusion

In summary, our study demonstrates the functional neural connections of the left amygdala formed during fear conditioning and subsequent extinction may be preserved in RS condition after several hours and days in healthy participants with low to moderate levels of anxiety. This finding implies that the left amygdala may play a more sustained role in long-term memory retention than the right amygdala in humans. Our results suggest the functional lateralization of the amygdala should be considered in further studies of fear memory and its pathophysiology in anxiety- and stress-related disorders.

## Declarations of interest

none

## Author Contributions

OM, AT, VB, GP, VU, and AM conceived and designed the study; AT and GP performed the experiments; VB, AT and OM analyzed the data; OM wrote the initial draft of the paper; all authors revised the article.

## List of Abbreviations

ACC: anterior cingulate cortex
BA: Brodmann area
BHISC: breath-holding induced signal changes
CS−: conditioned stimulus without unconditioned stimulus
CS+: conditioned stimulus with unconditioned stimulus
FC: functional connectivity
FE: fear extinction
FL: fear learning
FoV: field of view
FWHM: full width at half maximum
GLM: general linear model
HRF: hemodynamic response function
MNI: Montreal Neurological Institute space
MEG: magnetoencephalography
PFC: prefrontal cortex
PTSD: post-traumatic stress disorders
ROI: regions of interest
RS: resting state
rsFC: resting-state functional connectivity
SCR: skin conductance response
STAI: State-Trait Anxiety Inventory
US: unconditioned stimulus
VDM: voxel displacement map
WS: white screen.

## Funding

The Russian Scientific Foundation supported this work (Project No. 16-15-00300).

## Supporting Information

Supplementary data regarding this study can be found at https://www.dropbox.com/s/p8lcfuxohmlbb91/Supplementary%20Materials_v2.docx?dl=0

## Notes

https://www.dropbox.com/s/p8lcfuxohmlbb91/Supplementary%20Materials_v2.docx?dl=0

