## Supplemental Table S1 for "Longitudinal Changes of Resting-State Functional Connectivity of Amygdala Following Fear Learning and Extinction"

**Table S1.** Pseudorandom sequences of stimuli in the first session of fear learning (FL1), second session (FL2) and in the fear extinction (FE) session.

| | Fear Learning (FL1) | | | | | | Fear Learning 2 (FL2) | | | | | | Fear Extinction | | | | |
|---|---|---|---|---|---|---|---|---|---|---|---|---|---|---|---|---|---|
| № | Fixation cross (duration in s) | Type of CS | Duration of CS | US | Duration of white screen after CS | Total block duration | Fixation cross (duration in s) | Type of CS | Duration of CS | US | Duration of white screen after CS | Total block duration | Fixatio n cross (duratio n in s) | Type of CS | Duration of CS | Duration of white screen after CS | Total block duration |
| 1 | 2 | CS1+ | 8 | | 8 | 18 | 2 | CS- | 8 | | 8 | 18 | 2 | CS1+ | 6 | 10 | 18 |
| 2 | 2 | CS1+ | 8 | | 10 | 20 | 2 | CS- | 8 | | 10 | 20 | 2 | CS2+ | 8 | 8 | 18 |
| 3 | 2 | CS- | 8 | | 10 | 20 | 2 | CS1+ | 8 | | 10 | 20 | 2 | CS1+ | 6 | 10 | 18 |
| 4 | 2 | CS2+ | 6 | + | 10 | 18 | 2 | CS1+ | 6 | + | 10 | 18 | 2 | CS- | 6 | 12 | 20 |
| 5 | 2 | CS2+ | 8 | | 8 | 18 | 2 | CS1+ | 8 | | 8 | 18 | 2 | CS1+ | 8 | 12 | 22 |
| 6 | 2 | CS- | 4 | | 12 | 18 | 2 | CS- | 4 | | 12 | 18 | 2 | CS- | 6 | 12 | 20 |
| 7 | 2 | CS2+ | 6 | | 10 | 18 | 2 | CS2+ | 6 | | 10 | 18 | 2 | CS2+ | 4 | 10 | 16 |
| 8 | 2 | CS- | 4 | | 12 | 18 | 2 | CS2+ | 4 | + | 12 | 18 | 2 | CS- | 6 | 12 | 20 |
| 9 | 2 | CS2+ | 8 | | 8 | 18 | 2 | CS- | 8 | | 8 | 18 | 2 | CS- | 4 | 10 | 16 |
| 10 | 2 | CS1+ | 8 | + | 12 | 22 | 2 | CS2+ | 8 | + | 12 | 22 | 2 | CS1+ | 6 | 10 | 18 |
| 11 | 2 | CS2+ | 6 | | 12 | 20 | 2 | CS- | 6 | | 12 | 20 | 2 | CS2+ | 8 | 8 | 18 |
| 12 | 2 | CS2+ | 8 | | 10 | 20 | 2 | CS1+ | 8 | | 10 | 20 | 2 | CS- | 8 | 12 | 22 |
| 13 | 2 | CS1+ | 4 | + | 8 | 14 | 2 | CS- | 4 | | 8 | 14 | 2 | CS1+ | 4 | 10 | 16 |
| 14 | 2 | CS2+ | 6 | + | 10 | 18 | 2 | CS1+ | 6 | | 10 | 18 | 2 | CS2+ | 6 | 12 | 20 |
| 15 | 2 | CS- | 6 | | 8 | 16 | 2 | CS2+ | 6 | + | 8 | 16 | 2 | CS- | 4 | 10 | 16 |
| 16 | 2 | CS- | 6 | | 10 | 18 | 2 | CS1+ | 6 | + | 10 | 18 | 2 | CS1+ | 8 | 10 | 20 |
| 17 | 2 | CS2+ | 8 | | 10 | 20 | 2 | CS- | 8 | | 10 | 20 | 2 | CS2+ | 6 | 8 | 16 |
| 18 | 2 | CS1+ | 6 | + | 12 | 20 | 2 | CS- | 6 | | 12 | 20 | 2 | CS2+ | 6 | 10 | 18 |

| | | | | | | | | | | | | | | | | | |
|---|---|---|---|---|---|---|---|---|---|---|---|---|---|---|---|---|---|
| 19 | 2 | CS- | 4 | | 8 | 14 | 2 | CS2+ | 4 | | 8 | 14 | 2 | CS1+ | 8 | 8 | 18 |
| 20 | 2 | CS1+ | 6 | + | 12 | 20 | 2 | CS2+ | 6 | + | 12 | 20 | 2 | CS1+ | 6 | 10 | 18 |
| 21 | 2 | CS1+ | 6 | | 8 | 16 | 2 | CS1+ | 6 | | 8 | 16 | 2 | CS1+ | 8 | 8 | 18 |
| 22 | 2 | CS- | 8 | | 10 | 20 | 2 | CS- | 8 | | 10 | 20 | 2 | CS2+ | 6 | 10 | 18 |
| 23 | 2 | CS1+ | 6 | + | 8 | 16 | 2 | CS1+ | 6 | + | 8 | 16 | 2 | CS- | 6 | 12 | 20 |
| 24 | 2 | CS1+ | 4 | + | 10 | 16 | 2 | CS2+ | 4 | + | 10 | 16 | 2 | CS2+ | 8 | 10 | 20 |
| 25 | 2 | CS- | 8 | | 12 | 22 | 2 | CS- | 8 | | 12 | 22 | 2 | CS- | 4 | 10 | 16 |
| 26 | 2 | CS- | 4 | | 8 | 14 | 2 | CS2+ | 4 | + | 8 | 14 | 2 | CS2+ | 4 | 8 | 14 |
| 27 | 2 | CS1+ | 6 | + | 8 | 16 | 2 | CS1+ | 6 | | 8 | 16 | 2 | CS2+ | 4 | 12 | 18 |
| 28 | 2 | CS2+ | 4 | | 10 | 16 | 2 | CS1+ | 4 | | 10 | 16 | 2 | CS- | 6 | 8 | 16 |
| 29 | 2 | CS2+ | 4 | + | 8 | 14 | 2 | CS2+ | 4 | | 8 | 14 | 2 | CS2+ | 8 | 10 | 20 |
| 30 | 2 | CS- | 4 | | 10 | 16 | 2 | CS2+ | 4 | + | 10 | 16 | 2 | CS1+ | 8 | 8 | 18 |
| 31 | | | | | | | | | | | | | 2 | CS- | 6 | 12 | 20 |
| 32 | | | | | | | | | | | | | 2 | CS- | 6 | 10 | 18 |
| 33 | | | | | | | | | | | | | 2 | CS1+ | 4 | 10 | 16 |
| Total duration of session (s) | | | | | | 534 | | | | | | 534 | | | | | 600 |

CS1+, CS2+ - two types of conditioned stimuli with partial reinforcement, CS- one conditioned stimulus without reinforcement by unconditioned stimulus – an electrical shock
