## Supplemental Figure S1 for "Longitudinal Changes of Resting-State Functional Connectivity of Amygdala Following Fear Learning and Extinction"

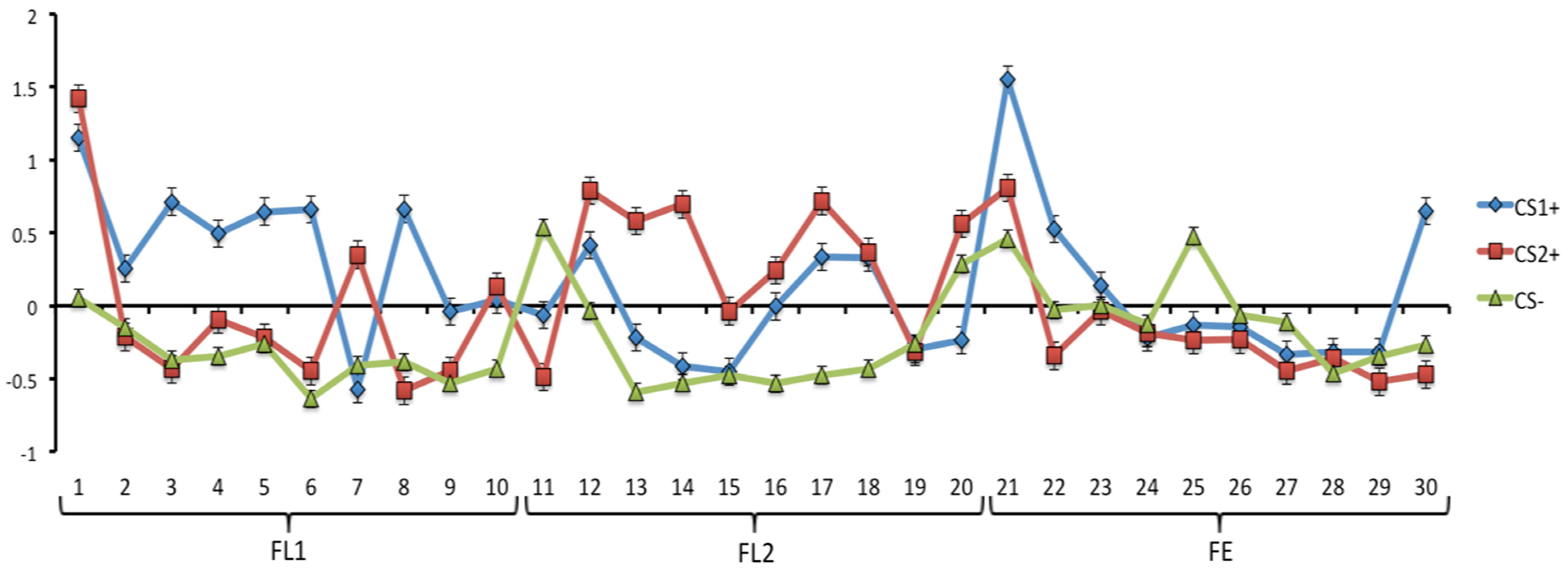


**Figure S1.** Skin conductance responses to the CS+ and CS-. Normalized skin conductance responses in the experimental group throughout the 10 CS1+, 10 CS2+ and 10 CS- trials in the three sessions: FL1- the first session of fear learning procedure, FL2- the second session of fear learning, and FE – fear extinction during fMRI.
