## Supplemental Table S2 for "Longitudinal Changes of Resting-State Functional Connectivity of Amygdala Following Fear Learning and Extinction"

**Table S2.** Brain areas exhibited a significant event-related BOLD response in the contrast of White screen after Conditioned Stimuli in the retention (A) and extinction stage (B) of fear in the experimental group (cluster size threshold > 169, $P_{FWE}<0.05$, $T<3.485$). R-in the right, L- in the left hemisphere (H); BA- Brodmann area, NoV – number of voxels.

| Anatomical region with peak intensity | H | BA | Peak MNI coordinate | | | t-value | NoV |
|---|---|---|---|---|---|---|---|
| | | | x | y | z | | |
| A -Retention | | | | | | | |
| Inferior Occipital Gyrus | L | 19 | -46.5 | -78 | -6 | 7.20 | 699 |
| Middle Occipital Gyrus | R | 19 | 49.5 | -78 | -6 | 6.88 | 1280 |
| Supramarginal Gyrus | R | 40 | 51 | -40.5 | 45 | 6.62 | 358 |
| Inferior Frontal Gyrus | L | 44 | -58.5 | 12 | 1.5 | 6.11 | 288 |
| Middle Frontal Gyrus | R | 9, 8 | 52.5 | 18 | 42 | 5.79 | 436 |
| Lingual Gyrus | L | 18 | -7.5 | -91.5 | -13.5 | 5.57 | 261 |
| Middle Frontal Gyrus | R | 10 | 43.5 | 57 | 13.5 | 5.06 | 180 |
| Middle Frontal Gyrus | L | 10 | -39 | 49.5 | 21 | 5.00 | 223 |
| Inferior Frontal Gyrus | R | 47 | 48 | 22.5 | 6 | 4.97 | 412 |
| Supramarginal Gyrus | L | 40 | -52.5 | -43.5 | 30 | 4.72 | 229 |
| Middle Occipital Gyrus | L | 18 | -22.5 | -94.5 | 9 | 4.60 | 250 |
| B - Extinction | | | | | | | |
| Lingual Gyrus | L | 19, 18 | -15 | -63 | -9 | 5.29 | 291 |
| Lingual Gyrus | L | 18 | -4.5 | -81 | 3 | 4.27 | 178 |
| Medial Frontal Gyrus | L | 10 | 3 | 66 | 4.5 | 6.0287 | 338 |
| Parahippocampal Gyrus | L | 30 | -16.5 | -54 | 1.5 | 5.8458 | 320 |
| Precuneus | R | 31 | 10.5 | -75 | 19.5 | 5.9981 | 1226 |
| Middle Occipital Gyrus | L | 19 | -45 | -78 | -7.5 | -6.43 | 647 |
| Middle Occipital Gyrus | R | 19 | 37.5 | -81 | -7.5 | -5.86 | 667 |
| Lingual Gyrus | R | 18 | 16.5 | -88.5 | -7.5 | -5.09 | 593 |
| Lingual Gyrus | L | 17 | -12 | -97.5 | -3 | -4.93 | 351 |
