## Supplemental Table S3 for "Longitudinal Changes of Resting-State Functional Connectivity of Amygdala Following Fear Learning and Extinction"

**Table S3.** Functional connectivity of amygdalar lateral seeds with other brain regions in the resting state immediately after fear extinction (A), in one day (B) and in 1 week (C) after fear learning and extinction: cluster threshold > 184 voxels, $P_{FWE}<0.05$). R-in the right, L- in the left hemisphere (H); BA- Brodmann area, NoV – number of voxels.

| Anatomical region with peak intensity | H | BA | Peak MNI coordinate | | | t-value | NoV |
|---|---|---|---|---|---|---|---|
| | | | x | y | z | | |
| A - Left Amygdala FC | | | | | | | |
| Precuneus | R | 7 | 1.5 | -54 | 36 | 8.82 | 209 |
| Supplementary Motor Area | R | 6 | 3 | 12 | 57 | 6.85 | 231 |
| Middle Occipital Gyrus | L | 19 | -46.5 | -85.5 | 9 | 6.70 | 345 |
| Precuneus | L | 7 | -9 | -57 | 72 | 6.50 | 676 |
| Middle Temporal Gyrus | L | 21 | -54 | -19.5 | -9 | 6.45 | 277 |
| Middle Frontal Gyrus | L | 9 | -27 | 28.5 | 39 | 6.43 | 472 |
| Superior Frontal Gyrus | L | 6 | -18 | 4.5 | 64.5 | 6.03 | 504 |
| Postcentral Gyrus | L | 2 | -24 | -37.5 | 76.5 | 5.91 | 487 |
| Superior Temporal Pole | L | 22 | -57 | 10.5 | -1.5 | 5.43 | 210 |
| Midcingulate Area | L | 5 | -4 | 43.5 | 48 | 5.42 | 408 |
| B - Left Amygdala FC: | | | | | | | |
| Middle Temporal Gyrus | R | 21 | 60 | -19.5 | -7.5 | 5.9329 | 203 |
| Cuneus | R | 17 | 6 | 97.5 | 0 | 5.6964 | 288 |
| Precentral Gyrus | L | 6 | 34.5 | -7.5 | 67.5 | 5.2214 | 331 |
| B - Right Amygdala FC: | | | | | | | |
| Inferior Frontal Gyrus, pars opercularis | R | 45 | 57 | 19.5 | 12 | 5.2534 | 306 |
| C - Left Amygdala FC: | | | | | | | |
| Middle Temporal Gyrus | R | 21 | 70.5 | -16.5 | -10.5 | 6.94 | 204 |
| Superior Temporal Gyrus | R | 21 | 45 | -7.5 | -12 | 6.92 | 266 |
| Superior Frontal Gyrus | L | 6 | -21 | 0 | 73.5 | 6.10 | 305 |
| Cingulate Gyrus | R | 31 | 19.5 | -40.5 | 36 | 5.95 | 328 |
| Superior Frontal Gyrus | R | 8 | 25.5 | 27 | 54 | 5.93 | 410 |
| Middle Temporal Gyrus | L | 19 | -43.5 | -85.5 | 21 | 5.72 | 417 |
| Calcarine Sulcus | R | 17 | 7.5 | -96 | 1.5 | 5.65 | 817 |
| Inferior Frontal Gyrus, pars orbitalis | R | 47 | 33 | 21 | -21 | 5.45 | 265 |
| Midcingulate Area | L | 31 | -4.5 | -43.5 | 48 | 5.39 | 386 |
| Middle Temporal Gyrus | R | 39 | 40.5 | -66 | 19.5 | 5.32 | 183 |
| Superior Temporal Pole | R | 38 | 55.5 | 9 | -12 | 4.48 | 236 |
| C - Right Amygdala FC: | | | | | | | |
| Middle Frontal Gyrus | R | 10 | 30 | 66 | 12 | 7.5103 | 790 |
| Inferior Frontal Gyrus, pars opercularis | R | 45 | 52.5 | 13.5 | 0 | 5.5058 | 548 |
| Superior Parietal Lobule | L | 7 | -36 | -60 | 60 | 5.2375 | 198 |
