## Supplemental Table S4 for "Longitudinal Changes of Resting-State Functional Connectivity of Amygdala Following Fear Learning and Extinction"

**Table S4.** Functional connectivity of the amygdalar lateral seeds with other brain regions in the resting state according to two-saple t-tests (Experimental group – Control group) for (A) the first resting state session before fear conditioning and (B) in one day after the first session (cluster threshold > 217 voxels, $P_{FWE}$<0.05), and C – in one week after fear learning and extinction (cluster threshold > 213 voxels, $P_{FWE}$<0.05). R-in the right, L- in the left hemisphere (H); BA- Brodmann area, NoV – number of voxels.

| Anatomical region with peak intensity | H | BA | Peak MNI coordinate | | | t-value | NoV |
|---|---|---|---|---|---|---|---|
| | | | x | y | z | | |
| A – Left Amygdala | | | | | | | |
| Middle Frontal Gyrus | L | 10 | -21 | 49.5 | 13.5 | -5.1966 | 238 |
| B- Left Amygdala FC | | | | | | | |
| Lingual Gyrus | L | 18 | -13.5 | -70.5 | -6 | 5.46 | 333 |
| Paracentral Lobule | L | 6 | -16.5 | -21 | 79.5 | 5.29 | 401 |
| Superior Occipital Gyrus | L | 18 | -13.5 | -90 | 18 | 4.8 | 242 |
| C - Left Amygdala FC | | | | | | | |
| Paracentral Lobule | L | 4 | -1.5 | -37.5 | 61.5 | 6.00 | 4218 |
| Precentral Gyrus | R | 6 | 48 | -12 | 43.5 | 5.88 | 895 |
| Precentral Gyrus | L | 6 | -54 | -6 | 46.5 | 5.01 | 383 |
| Calcarine Sulcus | R | 23 | 12 | -75 | 10.5 | 5.01 | 278 |
| Insula | R | 13 | 40.5 | -16.5 | 13.5 | 4.98 | 1861 |
| Midcingulate Area | R | 32 | 10.5 | 15 | 36 | 4.98 | 769 |
| Middle Temporal Gyrus | R | 39 | 51 | -72 | 13.5 | 4.82 | 244 |
| Superior Temporal Gyrus | R | 22 | 63 | -39 | 15 | 4.55 | 379 |
| Calcarine Sulcus | L | 17 | -3 | -79.5 | 9 | 4.49 | 372 |
| Cuneus | L | 7 | 1.5 | -72 | 33 | 4.07 | 280 |
| Superior Temporal Gyrus | L | 41 | -49.5 | -30 | 13.5 | 4.05 | 274 |
| C- Right Amygdala FC: | | | | | | | |
| Paracentral Lobule | R | 5 | 9 | -40.5 | 58.5 | 5.5255 | 963 |
