## Supplemental Table S5 for "Longitudinal Changes of Resting-State Functional Connectivity of Amygdala Following Fear Learning and Extinction"

**Table S5.** Areas of spatial overlap between cluster of activation in the retention stage of FE and differences in the amygdala rsFC between sessions.

| **Overlap with the event-relaed BOLD signal in the retention stage of FE** | | | | | | | | | | | | | | | |
|---|---|---|---|---|---|---|---|---|---|---|---|---|---|---|---|
| *Left Amygdala rsFC in RS_FE versus RS_0* | | | | | | | | *Right Amygdala rsFC in RS_FE versus RS_0* | | | | | | | |
| **NN** | **area** | **side** | **x** | **y** | **z** | **size** | **ba** | **NN** | **area** | **side** | **x** | **y** | **z** | **size** | **ba** |
| 1 | Middle Occipital Gyrus | L | -45,0 | -84,0 | -4,5 | 33 | 19 | | NaN | | | | | | |
| 2 | Rolandic Operculum | L | -55,5 | 15,0 | -4,5 | 63 | 22+4 | | | | | | | | |
| *Left Amygdala rsFC in RS_1 versus RS_0* | | | | | | | | *Right Amygdala rsFC in RS_1 versus RS_0* | | | | | | | |
| **NN** | **area** | **side** | **x** | **y** | **z** | **size** | **ba** | **NN** | **area** | **side** | **x** | **y** | **z** | **size** | **ba** |
| | NaN | | | | | | | 1 | Inferior Frontal Gyrus, pars orbitalis | R | 49,5 | 18 | -6 | 30 | 47 |
| *Left Amygdala rsFC in RS_7 versus RS_0* | | | | | | | | *Right Amygdala rsFC in RS_7 versus RS_0* | | | | | | | |
| **NN** | **area** | **side** | **x** | **y** | **z** | **size** | **ba** | **NN** | **area** | **side** | **x** | **y** | **z** | **size** | **ba** |
| 1 | Inferior Frontal Gyrus, pars orbitalis | R | 52,2 | 18 | -12 | 41 | 38+47 | 1 | Inferior Frontal Gyrus | R | 51 | 16,5 | -6 | 13 | 47 |
| | | | | | | | | 2 | Inferior Frontal Gyrus, pars opercularis | R | 58,5 | 18 | 0 | 5 | 47 |
| | | | | | | | | 3 | Middle Frontal Gyrus | R | 45 | 54 | 10,5 | 107 | 46+10 |
| **Overlap in rsFC between days** | | | | | | | | | | | | | | | |
| *Left Amygdala rsFC RS_FE versus RS_0 and RS_1 versus RS_0* | | | | | | | | *Right Amygdala rsFC RS_FE versus RS_0 and RS_1 versus RS_0* | | | | | | | |
| **NN** | **area** | **side** | **x** | **y** | **z** | **size** | **ba** | **NN** | **area** | **side** | **x** | **y** | **z** | **size** | **ba** |
| 1 | Superior Frontal Gyrus | L | -21 | 4,5 | 61,5 | 11 | wm | | NaN | | | | | | |
| *Left Amygdala rsFC RS_FE versus RS_0 and RS_7 versus RS_0* | | | | | | | | *Right Amygdala rsFC RS_FE versus RS_0 and RS_7 versus RS_0* | | | | | | | |
| **ba** | **area** | **side** | **x** | **y** | **z** | **size** | **ba** | **NN** | **area** | **side** | **x** | **y** | **z** | **size** | **ba** |
| 1 | Middle Temporal Gyrus | L | -44 | -87 | 16,5 | 16 | 19+39 | | NaN | | | | | | |
| 2 | Precuneus | L | -3 | -56 | 30 | 34 | 31+7 | | | | | | | | |
| 3 | Paracentral Lobule | L | -4,5 | -44 | 45 | 125 | 5+7 | | | | | | | | |
| 4 | Superior Frontal Gyrus | L | -18 | 0 | 69 | 33 | 6 | | | | | | | | |
| *Left Amygdala rsFC RS_1 versus RS_0 and RS_7 versus RS_0* | | | | | | | | *Right Amygdala rsFC RS_1 versus RS_0 and RS_7 versus RS_0* | | | | | | | |
| **NN** | **area** | **side** | **x** | **y** | **z** | **size** | **ba** | **NN** | **area** | **side** | **x** | **y** | **z** | **size** | **ba** |
| 1 | Lingual Gyrus | R | 10,5 | -95 | -17 | 216 | 17+18 | 1 | Inferior Frontal Gyrus | R | 51 | 15 | -4,5 | 31 | 47 |
| 2 | Middle Frontal Gyrus | L | -32 | 4,5 | 63 | 53 | 6 | 2 | Inferior Frontal Gyrus | R | 57 | 19,5 | 0 | 183 | 47+45+44 |
| 3 | Superior Frontal Gyrus | L | -32 | -7,5 | 66 | 115 | 6 | | | | | | | | |
