## Supplemental Table S6 for "Longitudinal Changes of Resting-State Functional Connectivity of Amygdala Following Fear Learning and Extinction"

**Table S6.** Correlation between state and trait anxiety and event-related BOLD response in fear extinction session, and functional connectivity of the left and right amygdala. R-in the right, L- in the left hemisphere (H); BA- Brodmann area, NoV – number of voxels.

| Anatomical region with peak intensity | H | BA | Peak MNI coordinate | | | t-value | NoV |
|---|---|---|---|---|---|---|---|
| | | | x | y | Z | | |
| **BOLD response and trait anxiety scores (k=174)** | | | | | | | |
| Middle Frontal Gyrus, orbital part | R | 11 | 30 | 45 | -10.5 | -5.1441 | 190 |
| Inferior Frontal Gyrus, pairs triangularis | L | 46 | -40.5 | 27 | 16.5 | -4.2637 | 182 |
| **Experimental group: RS_0 rsFC of the right amygdala with trait anxiety scores (k=160)** | | | | | | | |
| Medial Frontal Gyrus | R | 10 | 9 | 66 | 3 | 6.4297 | 246 |
| **Control group: RS_1 rsFC of the left amygdala with trait anxiety scores (k=176)** | | | | | | | |
| Hippocampus | R | - | 37.5 | -28.5 | -10.5 | 7.5946 | 222 |
| **Control group: RS_7 rsFC of the right amygdala with state anxiety scores (k=162)** | | | | | | | |
| Precuneus | R | 7 | 18 | -66 | 36 | 6.3738 | 275 |
| Middle Frontal Gyrus | L | 6 | -33 | 9 | 63 | -5.8187 | 193 |
